# Localized heterochrony integrates overgrowth potential of oncogenic clones

**DOI:** 10.1101/2022.07.15.500211

**Authors:** Nicola Blum, Matthew P. Harris

## Abstract

Somatic oncogenic mutations are frequent and can occur early during development. The result is the formation of a patchwork of mutant clones. Such mosaicism has been implicated in a broad range of developmental anomalies however their etiology is poorly understood. Patients carrying a common somatic oncogenic mutation in either *PIK3CA* or *AKT1*, can present with disproportionally large digits or limbs. How mutant clones, carrying an oncogenic mutation that often drives unchecked proliferation leads to controlled and coordinated overgrowth is unknown. We use the zebrafish to explore growth dynamics of oncogenic clones during development. In subset of clones, we observe a local increase in proportion of the fin skeleton closely resembling patient overgrowth phenotypes. We unravel the cellular and developmental mechanisms of these overgrowths and pinpoint the cell type and timing of clonal expansion. Coordinated overgrowth is associated with rapid clone expansion during early pre-chondrogenic phase of bone development inducing a heterochronic shift that drives the change in bone size. Our study details how development integrates and translates growth potential of oncogenic clones, thereby shaping the phenotypic consequences of somatic mutations.

## INTRODUCTION

With the advent of whole genome sequencing of somatic tissues and cellular bar coding, it is becoming clear that somatic mutations are quite common (Biesecker and Spinner, 2013; García-Nieto et al., 2019; Li et al., 2021). Starting in the early embryo, somatic mutations accumulate in our cells throughout life leading to a patchwork of mutant cell clones in our tissues and organs. Among mutant clones a small minority may harbor a mutation that increases the growth potential of a cell. Such clones have long been known to cause cancer and it is becoming increasingly evident that a broad range of developmental disorders are also products of somatic oncogenic mutations (Moog et al., 2020; Mustjoki and Young, 2021; Nussinov et al., 2022). The underlying mechanisms determining the particular expressivity or presentation of oncogenic clones in development, however, remain largely unknown.

PROS (**P**IK3CA, the catalytic subunit of the phosphoinositide 3 kinase (PI3K), **R**elated **O**vergrowth **S**pectrum) and Proteus syndrome are extremely rare disorders characterized by localized overgrowths that can affect virtually any tissue and organ (Doucet et al., 2016; Keppler-Noreuil et al., 2014; Keppler-Noreuil et al., 2015; Kurek et al., 2012; Lindhurst et al., 2011; Marjorie and Lindhurst, 2019; Venot et al., 2018). Somatic activating mutations in PIK3CA (H1047R, E542K, E545K, G1049R) and AKT Serine/Threonine Kinase 1 (AKT1^E17K^) have been detected in patients with PROS and Proteus syndrome, respectively. These mutations are well-known oncogenic mutations (Chen et al., 2018; Karakas et al., 2006; Lawrence et al.; Rudolph et al., 2016; Shoji et al., 2009) that render the PI3K/AKT pathway, a key regulator of cell growth, proliferation and survival, hyperactive (Hoxhaj and Manning, 2020). In both PROS and Proteus syndromes, the particular mutation does not define the particular presentation as the same mutations are found to drive a suite of different disorders. PROS and Proteus syndrome are remarkably heterogeneous disorders. Phenotypes arising from these mutations are likely the result of modifying genetic factors or, alternatively, with the timing and location of mutation event in the embryo likely critically influencing phenotypic manifestation. The underlying mechanisms detailing the particular phenotypic expressivity of clones with oncogenic PI3K/AKT signaling, however, remain largely unknown.

Overgrowth in PROS and Proteus syndrome patients is often uncoordinated disrupting normal tissue patterning and proportions. Interesting exceptions however are cases where patients present with localized gigantism in the appendicular skeleton due to disproportionally large bones (**Fig. 1A, B**) (Bornstein et al., 2014; Cerrato et al., 2013; Keppler-Noreuil et al., 2014; Keppler-Noreuil et al., 2015; Rios et al., 2013; Tian et al., 2020; Tripolszki et al., 2016; Wu et al.; Zeng et al., 2020). In such patients, affected long bones are longer and wider, yet overall retain their normal shape indicating that growth is controlled and coordinated. The result is an abnormally large limb or digit (macrodactyly). In the latter case, often more than one, but always adjacent, digits are affected suggesting regional affected territories. How somatic oncogenic mutations in the PI3K/AKT pathway, which are associated with unchecked proliferation and a broad range of neoplasms, can lead to controlled and coordinated overgrowth of entire entities provides a useful case study to address how oncogenic potential is shaped in development. Here, we develop a zebrafish model for overgrowth observed in PROS and Proteus syndrome and explore the growth dynamics of mutant clones during bone development. Our findings reveal that over-proliferation of mutant clones is restricted to the pre-chondrogenic phase during bone development. Interestingly, the rapid expansion of mutant clones during this early stage causes premature skeletal condensation and this shift in developmental timing drives the observed non-neoplastic overgrowth of entire long bones.

**Figure 1.**
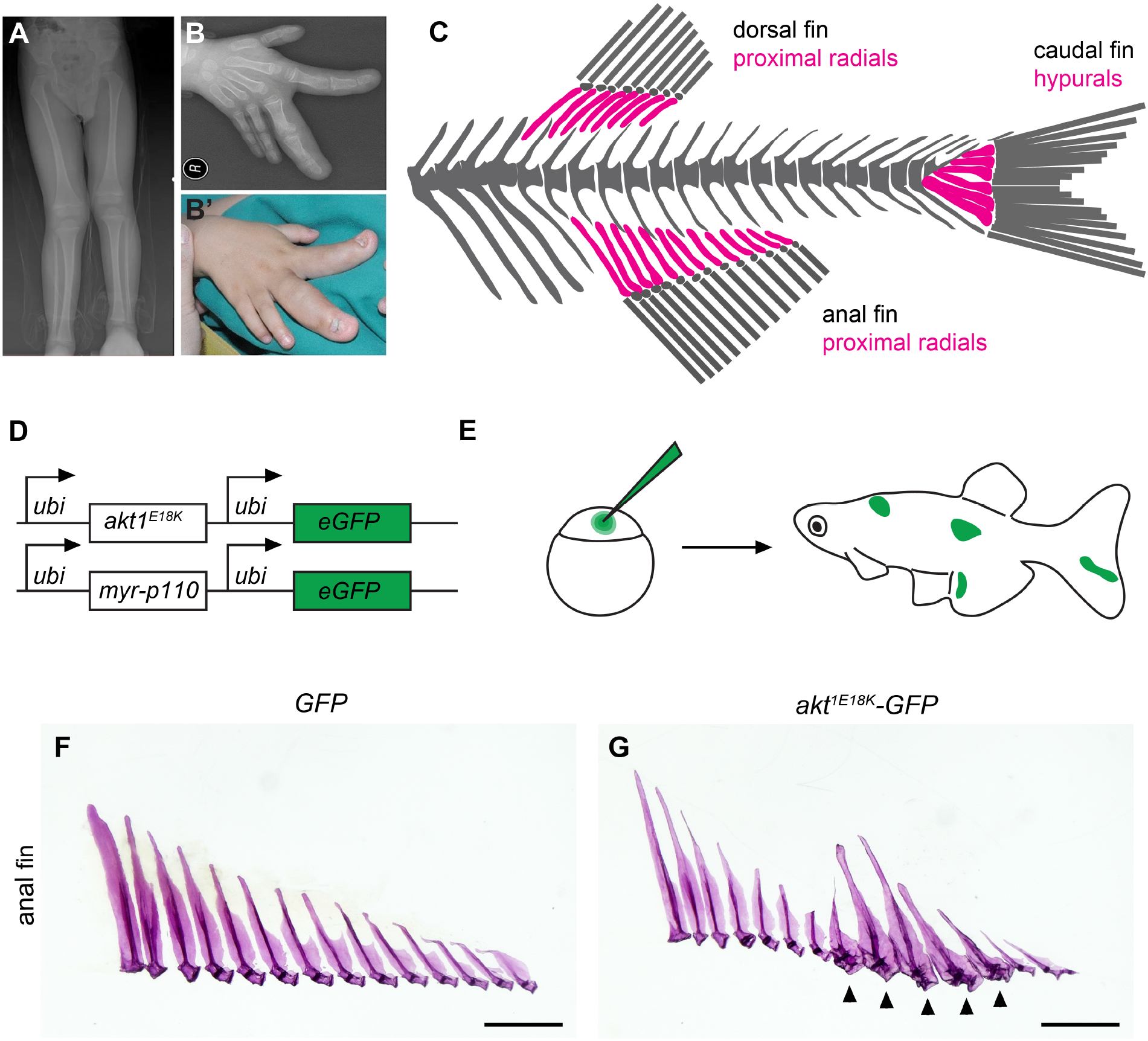
Mosaic overactivation of PI3K/AKT is sufficient to cause fin long bone overgrowth. (**A**) Leg length discrepancy due to overgrowth of the right tibia and fibula in a patient with Proteus syndrome. (**B**) Macrodactyly in a patient with PROS. (**C**) Generalized diagram of the adult zebrafish axial skeleton. Long bones of the fin endoskeleton are highlighted in magenta. (**D, E**) Experimental strategy to generate zebrafish harboring clones with oncogenic PI3K/AKT signaling. (**D**) Constructs used to create mosaic zebrafish via Tol2 transgenesis (**E**). (**F, G**) Mosaic overactivation of PI3K/AKT results in localized gigantism of the adult fin endoskeleton. Alizarin red stained preparations of anal fin proximal radials in control (**F**) and *akt1^E18K^-GFP* (**G**) mosaic fish. Arrowheads in (**G**) mark overgrown bones. Scale bars (**F, G**) 0.1 cm. Images in (**A, B**) are modified from Zeng et al. (Zeng et al., 2020) (**A**) and Tian et al. (Tian et al., 2020) (**B**) and licensed under CC by 4.0 (http://creativecommons.org/licenses/by/4.0/).

## RESULTS AND DISCUSSION

### Mosaic overactivation of PI3K/AKT signaling causes localized gigantism of the fin endoskeleton

To understand how mutant clones, harboring an oncogenic mutation in either PIK3CA or AKT1, can cause controlled overgrowth of long bones, we thought to employ a model that would allow live imaging of mutant cell clones. To this end we made use of zebrafish as we can readily make clones and observe them in real time over development and into adulthood (Perathoner et al., 2014). The endoskeleton of zebrafish median fins (caudal, anal and dorsal fin) harbors a series of long endochondral bones (**Fig. 1C**), called hypurals in the caudal fin and proximal radials in dorsal and anal fins (Bird and Mabee, 2003). Of note, hypurals and proximal radials have characteristic size ratios (*e.g*. in the anal fin, bone length decreases from anterior to posterior). Thus, overgrowth of individual bones can be readily visible. The cellular and molecular mechanisms underlying patterning and growth of long bones in the teleost fin are very similar to those in the tetrapod limb (Crotwell and Mabee, 2007; Hawkins et al., 2021).

To generate randomly distributed clones in embryos with overactivated PI3K/AKT signaling, we used a Tol2 transposase to allow random, mosaic stable transgene integration during early embryonic development (**Fig. 1D, E**) (Kawakami, 2007). Transgenic constructs harbor either *akt1^E18K^* (the zebrafish corresponding mutation of human *akt1^E17K^*) or a constitutively active form of murine PIK3CA (*myr-p110α*) under the control of the ubiquitously-active *ubiquitin (ubi*) promoter. Both constructs have a fluorescent marker (*eGFP*) under the control of a second *ubi* promoter to track the presence of clones. Hereafter, we refer to the transgenic constructs as *akt1^E18K^-GFP, myr-p110α-GFP* and *GFP* (control).

We found that high clone numbers of *akt1^E18K^-GFP* and *myr-p110α-GFP* lead to early lethality at larval stages due to severe vascular abnormalities and hemorrhaging (not shown). This is in congruence with findings in PROS models in the mouse (Venot et al., 2018). Vascular abnormalities are also common in patients with PROS or Proteus syndrome (Castillo et al., 2016; Detter et al., 2018; Luks et al., 2015; Perry et al., 2007; Queisser et al., 2018; Rodriguez-Laguna et al., 2019). To accommodate observation of clones during later development, we titrated clone numbers in such a way to minimize lethality. Even in these experimental conditions approximately 50% of injected fish harbored clones in multiple tissues. To verify that *akt1^E18K^-GFP* and *myr-p110α-GFP* mosaic animals mimic overgrowth phenotypes seen in PROS and Proteus syndrome patients, we analyzed adult fish (3-6 months old) for externally visible overgrowth phenotypes. Indeed, after injection at lower titrations, fish showed localized or patchy overgrowth in the skin, adipose tissue, muscle and vasculature (**Supplementary Fig 1A-C**).

We next tested whether *akt1^E18K^-GFP* and *myr-p110α-GFP* clones cause localized gigantism in the fin endoskeleton by analyzing alizarin red stained skeletons of mosaic fish having identified clones in adult fins. We found disproportionally large long bones in median fins: caudal (GFP, n=0; *akt1^E18K^-GFP*, n=7; *myr-p110α-GFP*, n=5), anal (GFP, n=0; *akt1^E18K^-GFP*, n=13; *myr-p110α-GFP*, n=8) and dorsal (GFP, n=0; *akt1^E18K^-GFP*, n=11; *myr-p110α-GFP*, n=5) fins (**Fig. 1F,G** and not shown). In most cases more than one, but always adjacent bones were affected. These structures closely resemble long bone overgrowth observed in patients with PROS and Proteus syndrome revealing an actuatable model for this disorder in the zebrafish.

### Overgrowth is caused by a heterochronic shift in development of the cartilage anlagen

To determine in which cell type the mutation has to occur in order cause long bone overgrowth, we generated *akt1^E18K^-GFP* and *myr-p110α-GFP* clones in transgenic lines labeling either chondrocytes (*Tg(sox10:DsRed)*) (Das and Crump, 2012) or osteoblasts (*Tg(Ola.Sp7:mCherry)*) (DeLaurier et al., 2010) to visualize the fin endoskeleton in live larval and juvenile fish. Imaging at 8-9 mm standard length (SL) revealed that long bone overgrowth is already present at late larval stages in caudal (GFP, n=0; *akt1^E18K^-GFP*, n=50; *myr-p110α-GFP*, n=39), anal (GFP, n=0; *akt1^E18K^-GFP*, n=63; *myr-p110α-GFP*, n=42) and dorsal (GFP, n=0; *akt1^E18K^-GFP*, n=72; *myr-p110α-GFP*, n=38) fins (**Fig. 2A, B** and not shown). In all overgrown bones, *akt1^E18K^-GFP* and *myr-p110α-GFP* colocalized with chondrocyte and osteoblast markers (**Fig. 2C, D** and not shown) as well as the surrounding connective tissue (**Fig. 2C, D**). Thus, the change in relative long bone size is caused by PI3K/AKT overactivation in the mesenchymal lineage.

**Figure 2.**
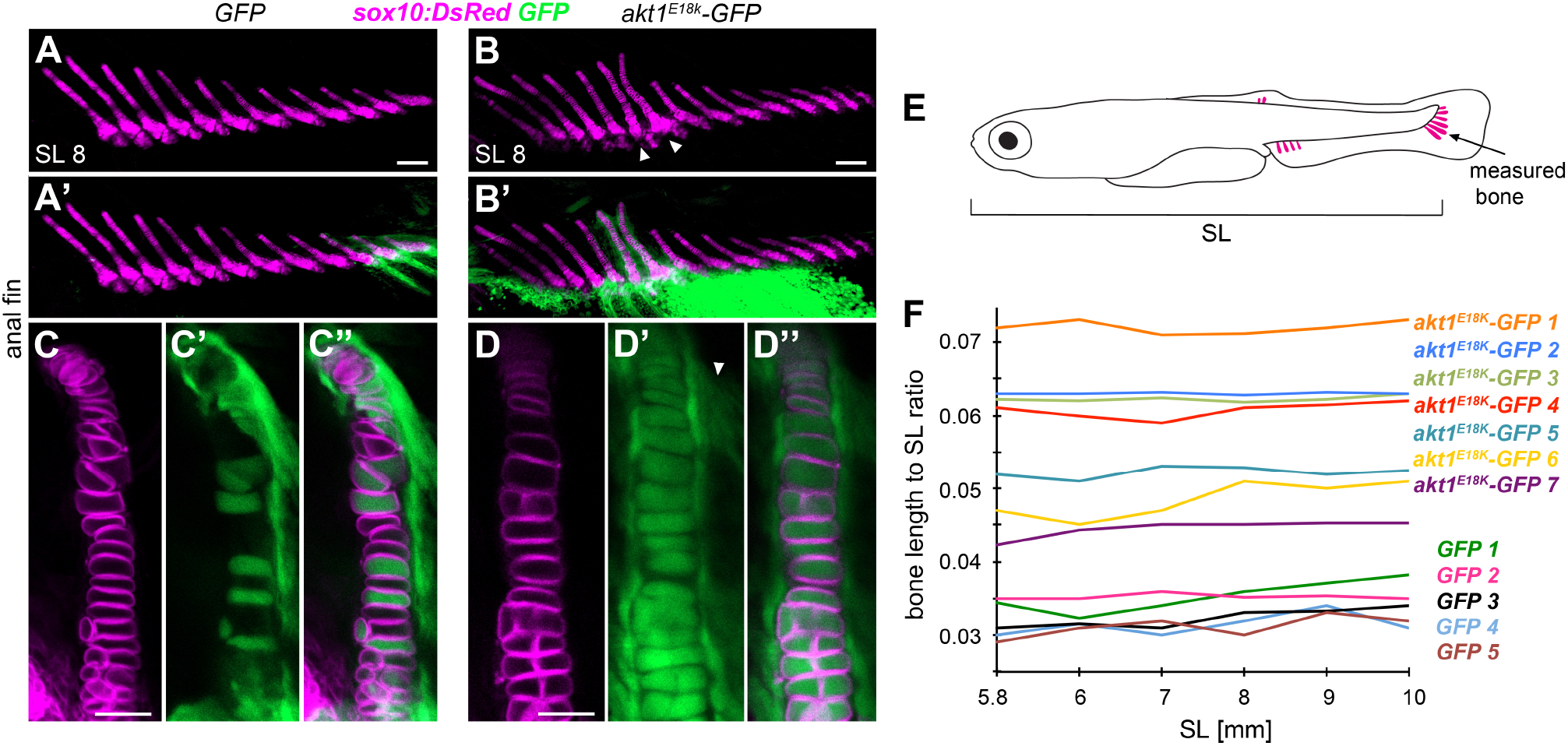
Bone overgrowth is nonprogressive and caused by PI3K/AKT overactivation in mesenchymal derivatives. (**A, B**) PI3K/AKT mosaic overactivation results in an early overgrowth phenotype in the fin endoskeleton. Live imaging of chondrocytes (marked by *Tg(sox10:DsRed)*) at SL 8 mm reveals localized skeletal overgrowth in the anal fin of an *akt1^E18K^-GFP* mosaic larva overlapping with GFP^+^ cells. (**A**) Control. (**B**) *akt1^E18K^-GFP*. 3D projections from z-stacks. Arrowheads in (**B**) mark affected bones. (**C, D**) Higher magnifications show that *akt1^E18K^-GFP* colocalizes with chondrocytes and adjacent fibroblasts (arrowhead in **D**’). (**C**) Control. (**D**) *akt1^E18K^-GFP*. Note that all chondrocytes in *akt1^E18K^-GFP* mosaic larvae harbor the transgene. (**E, F**) The change in relative long bone size is static in *akt1^E18K^-GFP* mosaic fish. (**E**) Analyzed bone (hypural 1 of the caudal fin). (**F**) Length of hypural 1 to SL ratio over time in *akt1^E18K^-GFP* (n=7) and *GFP (n=5*) mosaic larvae. Scale bars (**A, B**) 100 μm, (**C, D**) 25 μm.

We next explored the developmental mechanisms by which *akt1^E18K^-GFP* and *myr-p110α-GFP* clones modify bone size. A change in relative size can be achieved through three different ways: 1) a change in the rate of long bone elongation, 2) a change in developmental timing (heterochrony) or 3) a change in the size of cartilage anlagen. Similar to tetrapod long bones, elongation of long bones in fins is driven by chondrocyte proliferation and subsequent hypertrophy in growth zones at the end of bones. The PI3K/AKT pathway is a key regulator in promoting cell growth and proliferation suggesting that *akt1^E18K^-GFP* and *myr-p110α-GFP* positive chondrocytes may possess an increased growth potential. It is, therefore tempting to speculate that the observed change in bone size is due to accelerated bone elongation. Indeed, we found that in contrast to control larvae, all chondrocytes in affected long bones in *akt1^E18K^-GFP* and*myr-p110α-GFP* mosaic larva harbored the transgene (GFP n=0, *akt1^E18K^-GFP* n=53, *myr-p110α-GFP* n=34) (**Fig. 2C, D**) indicative for a growth advantage of affected cells over those of wild type. Surprisingly, we found that overgrowth is not progressive and affected bones grew at the same rate as bones in control larvae (**Fig 2E, F**). Thus, the observed change in relative long bone size is not caused by an increase in growth rate. Moreover, this finding demonstrates that over-proliferation of PI3K/AKT mutant clones is restricted to early bone development.

Next, we analyzed whether bone overgrowth is caused by a change in relative developmental timing. Using uninjected *Tg(sox10:DsRed*) fish, we found that the first cartilage anlagen appear at approximately SL 4.5 mm in the caudal fin (n= 17), at SL 5.5 mm in the anal fin (n= 15) and at SL 5.8 mm in the dorsal fin (n= 19) (**Fig. 3A-C**). Surprisingly, in *akt1^E18K^-GFP* and *myr-p110α-GFP* mosaic fish, cartilage anlagen with overlapping mesenchymal clones appeared prematurely as early as SL 4.0 mm in caudal (GFP n=0, *akt1^E18K^-GFP* n=93, *myr-p110α-GFP* n=45), anal (GFP n=0, *akt1^E18K^-GFP* n=85, *myr-p110α-GFP* n=51) and dorsal (GFP n=0, *akt1^E18K^-GFP* n=87, *myr-p110α-GFP* n=39) fins (**Fig. 3D-I** and not shown). Taken together, these findings demonstrate that mesenchymal clones, carrying an oncogenic mutation in the PI3K/AKT pathway cause a heterochronic shift in formation of cartilage anlagen and that this temporal modulation underlies the change in relative long bone size.

**Figure 3.**
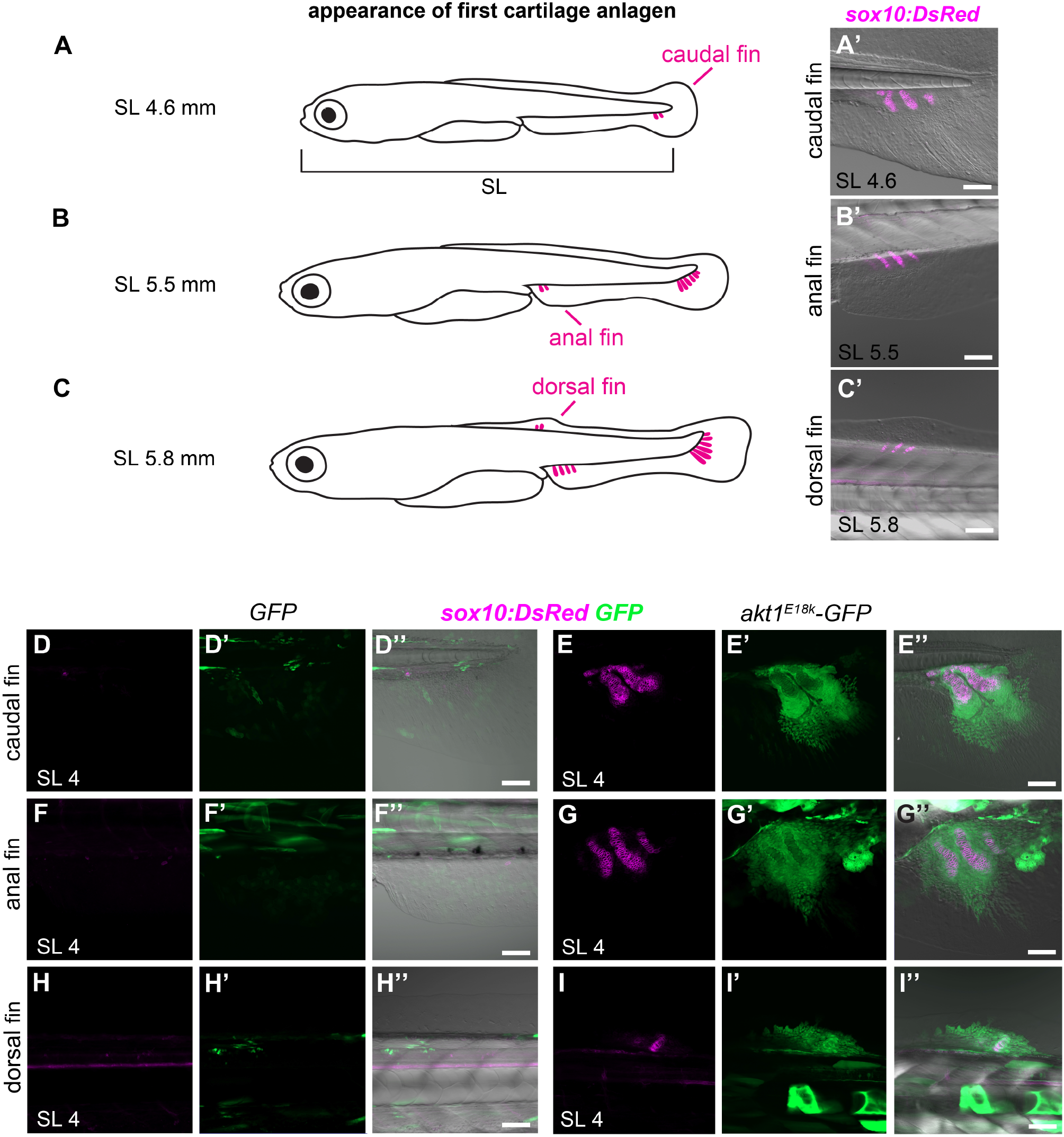
PI3K/AKT mutant clones induce a heterochronic shift in bone development. (**A-C**) Appearance of first cartilage anlagen in the caudal, anal and dorsal fin. (**A’, B’, C’**) Live imaging of first cartilage anlagen (marked by *Tg(sox10:DsRed)*) in caudal (**A’**), anal (**B’**) and dorsal (**C’**) fin. (**D-I**) *akt1^E18K^-GFP* mesenchymal clones induce premature formation of cartilage anlagen in caudal (**D, E**), anal (**F, G**) and dorsal (**H, I**) fins. (**D, F, H**) Control. (**E, G, I**) *akt1^E18K^-GFP*. Scale bars, 50 μm.

### PI3K/AKT mutant clones form premature pre-cartilage condensations

To explore how PI3K/AKT mutant mesenchymal clones cause the observed heterochronic shift in cartilage formation, we examined the behavior of mutant clones prior to the appearance of cartilage anlagen in development of the fins. Development of cartilage anlagen is initiated as undifferentiated, and seemingly homogenous pre-chondrogenic, mesenchymal cells become closely juxtaposed forming a pre-chondrogenic condensation. Subsequently, these condensations initiate matrix deposition and differentiate into chondrocytes (Hall et al., 1996). During the condensation process, critical cell-cell and cell-matrix interactions occur that are necessary to trigger chondrogenic differentiation. In zebrafish, pre-chondrogenic condensations of median fins are visible as thickenings in the fin fold and growth proceeds in a proximal to distal direction (Parichy et al., 2009).

We found that at these early stages of fin development, *akt1^E18K^-GFP* and *myr-p110α-GFP* mesenchymal clones were much larger than control clones (**Fig. 4A, B** and not shown) indicative of overproliferation of mutant clones. Notably, we did not detect large clones in adjacent trunk regions (**Fig. 4B**) demonstrating that expansion of *akt1^E18K^-GFP* and *myr-p110α-GFP* clones occurred after recruitment of progenitors into the fin. This finding is important for understanding the timing of mutation event in patients and suggests that the mutation can occur relatively late during embryogenesis, while still resulting in a high number of mutant cells prior cartilage formation. Intriguingly, *akt1^E18K^-GFP* and *myr-p110α-GFP* mutant clones formed premature pre-chondrogenic condensations in caudal (GFP n=0, *akt1^E18K^-GFP* n=101, *myr-p110α-GFP* n=45), anal (GFP n=0, *akt1^E18K^-GFP* n=90, *myr-p110α-GFP* n=34) and dorsal (GFP n=0, *akt1^E18K^-GFP* n=93, *myr-p110α-GFP* n=35) fins (**Fig. 4B** and not shown). We conclude that mesenchymal clones, carrying an oncogenic mutation in the PI3K/AKT pathway cause a heterochronic shift in bone formation through premature pre-chondrogenic condensation. This shift occurs during early bone development and later clones during bone elongation do not show comparable overgrowth resulting in proportional, or coordinated changes in size.

**Figure 4.**
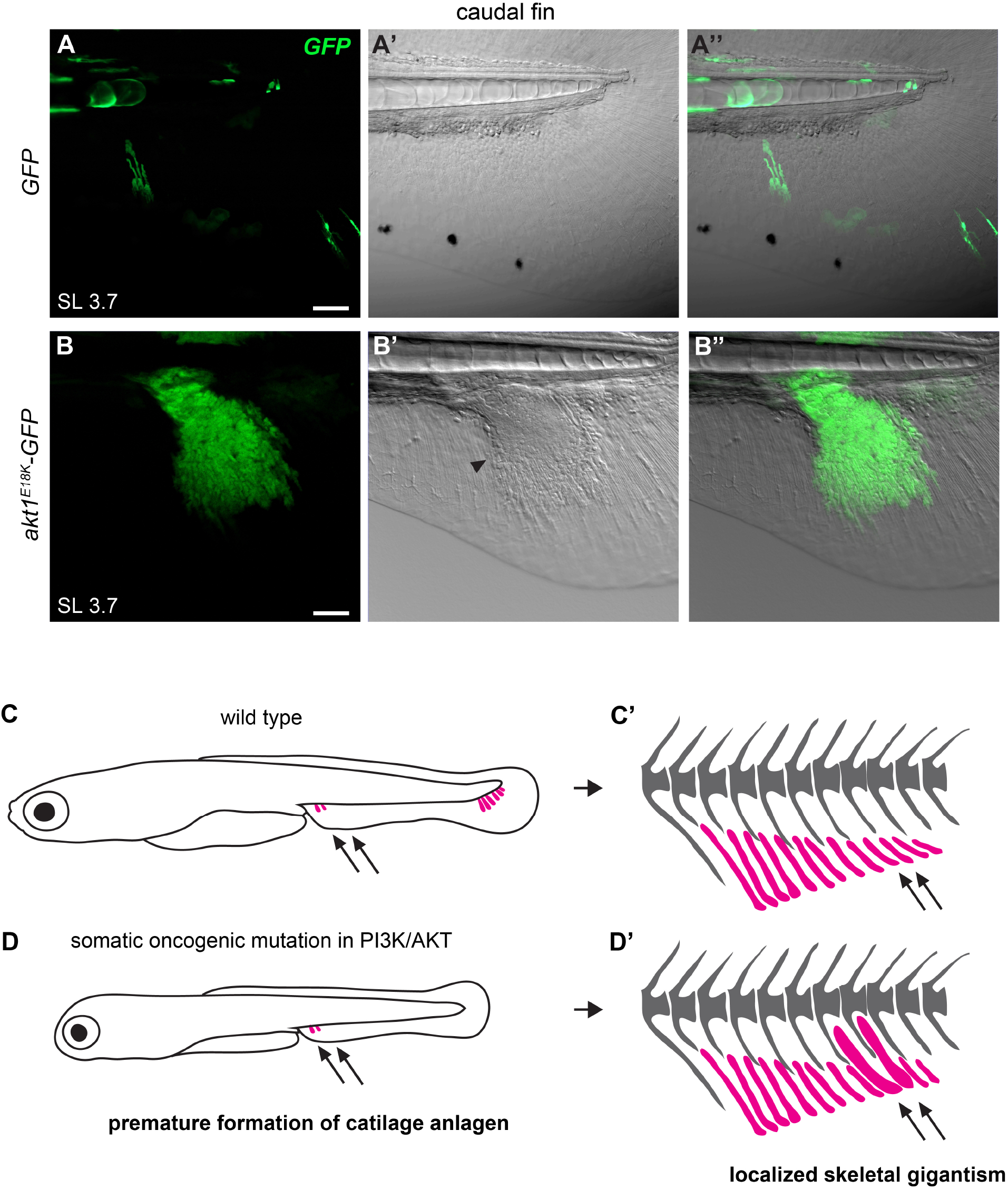
PI3K/AKT mutant mesenchymal clones form premature pre-chondrogenic condensations. **A, B**) Live imaging of *akt1^E18K^-GFP* mesenchymal clones at SL 3.7 mm reveals clonal expansion and premature pre-chondrogenic condensation in the caudal fin. Arrowhead in (**B**’) marks the edge of condensation. (**A**) Control. (**B**) *akt1^E18K^-GFP*. Scale bars, 50 μm. (**C, D**) Developmental mechanism underlying localized skeletal gigantism caused by somatic oncogenic mutations in the PI3K/AKT pathway.

Pre-chondrogenic condensation is dependent on cell number (Osdoby and Caplan, 1979; Osdoby and Caplan, 1980; San Antonio and Tuan, 1986). The burst in proliferation of PI3K/AKT oncogenic clones likely accelerates an increase in cell density prior to pre-chondrogenic condensation. As a result of this, the critical threshold in cell number is reached earlier. However, although high density of pre-chondrogenic cells is a requirement, it is not likely to be sufficient to induce condensation. Interestingly, mTor signaling, a primary mediator of PI3K/AKT signaling has previously been shown to be required for pre-chondrogenic condensation through translational control of SOX9 (Iezaki et al., 2018). We propose a model in which PI3K/AKT mutant clones induce premature condensation through both increasing cell density and upregulation of SOX9 levels.

### Conclusion

Somatic mutations are increasingly implicated as the underlying cause of many developmental disorders. However, we know very little in how developmental programs affect the behavior and phenotypic outcome of mutant clones, and further if these attributes can be leveraged to design interventions or treatments. By using a zebrafish model, we unraveled how overproliferation of oncogenic clones can lead to non-neoplastic and coordinated overgrowth of entire bones. We show that during the earliest phase of bone development, the growth potential of oncogenic clones is translated into a heterochronic shift in cartilage formation leading to localized, coordinated gigantism of the appendicular skeleton. Our findings support a model in which pre-chondrogenic cells carrying a somatic oncogenic mutation in *PIK3CA* or *AKT1* initiate cartilage development prematurely driving a qualitative change in relative long bone size (**Fig. 4C, D**); importantly, overgrowth in this context is not neoplastic, rather coordinated demonstrating a capacity within the developmental system for growth integration in these disorders.

Heterochrony is a pervasive developmental mechanism in evolution, underlying shifts in morphology and physiology (Beer, 1930; Gould, 1977). Through changes in developmental timing, broad changes can occur while remaining integrated and patterned. The regulation of differential timing of growth underlying heterochrony is generally unknown, but is most likely multifaceted. Our work shows a particular case in which qualitative shifts in size are modulated within a distinct window of the pre-chondrogenic phase of skeletal anlagen formation. Our work modeled localized, static overgrowth of long bones independent of other PROS and Proteus syndrome clinical presentations. However, patients with PIK3CA or AKT1 driven macrodactyly or enlarged skeletal bones can present with either static or progressive overgrowth (Cerrato et al., 2013; Keppler-Noreuil et al., 2014; Keppler-Noreuil et al., 2015). Therefore, heterochrony is only a component of the developmental modulation of oncogenic clone behavior in these disorders. It remains to be shown whether the differential growth dynamics of static or progressive growth is due to modulating factors or, alternatively independent mechanisms such as PI3K/AKT overactivation in a different cell type.

## MATERIAL AND METHODS

### Zebrafish husbandry, strains and staging

The study was conducted with ethical approval from the Institutional Animal Care and Use Committee of Boston Children’s Hospital. Zebrafish were maintained under standard conditions (WESTERFIELD, 2000). A description of the husbandry and environmental conditions is available at www.protocols.io/(dx.doi.org/10.17504/protocols.io.mrjc54n). All experiments were performed in the casper mutant background (White et al., 2008) to ensure optical accessibility of postembryonic stages. The following transgenic zebrafish strains were used: *Tg(sox10:DsRed)^el10Tg^* (Das and Crump, 2012) and *Tg(Ola.Sp7:mCherry)^zf131^* (DeLaurier et al., 2010). Standard length (SL) were used for staging of larvae and juvenile fish according to Parichy et al. (Parichy et al., 2009).

### Generation of mosaic zebrafish

The following plasmids were constructed for injection: *pmTol2-ubi:eGFP* (control construct), *pmTol2-ubi:akt^1E18K^-ubi:eGPF, pmTol2-ubi:myr-p110-ubi:eGPF. pminiTol2* (Addgene #31829) was used as backbone vector and transgene cassettes (*ubi:eGFP, ubi:akt1^E18K^* and *ubi:myr-p110*) were inserted into the multiple cloning site. Transgene cassettes were generated by inserting *eGFP* from *pME-eGFP* (Tol2kit #383), murine *myr-p110* from Myr-(iSH2-p85)-p110alpha-Myc (Addgene #1410) or zebrafish *akt1^E18K^* together with the SV40 late polyadenylation signal (SV40pA) downstream of the ubi promoter into *pENTR5’ubi* (Addgene #27320). The full length coding sequence of zebrafish *akt1* was amplified from 24 hours post fertilization cDNA and the E18K mutation was introduced using QuickChange Site-Directed mutagenesis (Agilent). To generate mosaic fish, plasmid DNA (5 ng/μl) and Tol2 transposase mRNA(4 ng/μl) (synthesized from *pT3TS-Tol2* Addgene #31831) were injected into one-cell stage embryos. Optimal DNA and Tol2 mRNA concentration were determined by titration experiments.

### Skeletal staining

Adult fish were fixed for 1-2 days in 3.7% formaldehyde in phosphate buffered saline (PBS). After fixation, fish were briefly rinsed with PBS and gradually transferred to 95% ethanol. Subsequently, fish were washed with acetone for 6 hours to remove fat. Fish were then briefly rinsed with 95% ETOH and gradually transferred to PBS. Fish were equilibrated in 0.5% potassium hydroxide (KOH) and stained for 24 hours in 0.1% Alizarin Red S (Sigma) in 0.5% KOH. After staining, fish were rinsed with 0.5% KOH followed by PBS. Next, fish were macerated using 3% trypsin (Fisher Scientific) in 30% saturated sodium borate at 37°C for 2-4 hours. Digestion was stopped by briefly rinsing fish with PBS. Subsequently, fish were further cleared in 2% KOH for 24 hours and gradually transferred to 75% glycerol in 0.5% KOH for storage and imaging. If not otherwise stated, all steps were carried out at room temperature with gentle rocking agitation.

### Imaging

Stained fish skeletons were imaged using Nikon SMZ18 stereomicroscope and NIS-Elements software. Adult fish were anesthetized with MS-222 and photographed using a Canon EOS digital camera. Live imaging of larval fish were performed using a Zeiss LSM 800 confocal microscope and ZEN Blue software. Larvae were anesthetized with MS-222 and embedded in 0.5% low melting agarose in E3 for imaging. Confocal z-stacks were processed using ZEN blue software.

## ACKNOWLEDGEMENTS

This work was partially supported by NIH R01HD084985 and BSF US-Israel BSF (grant no. 2017204) to MPH as well as a DFG research fellowship (BL1614/1-1) to NB.

---

**Supplementary Figure 1.**
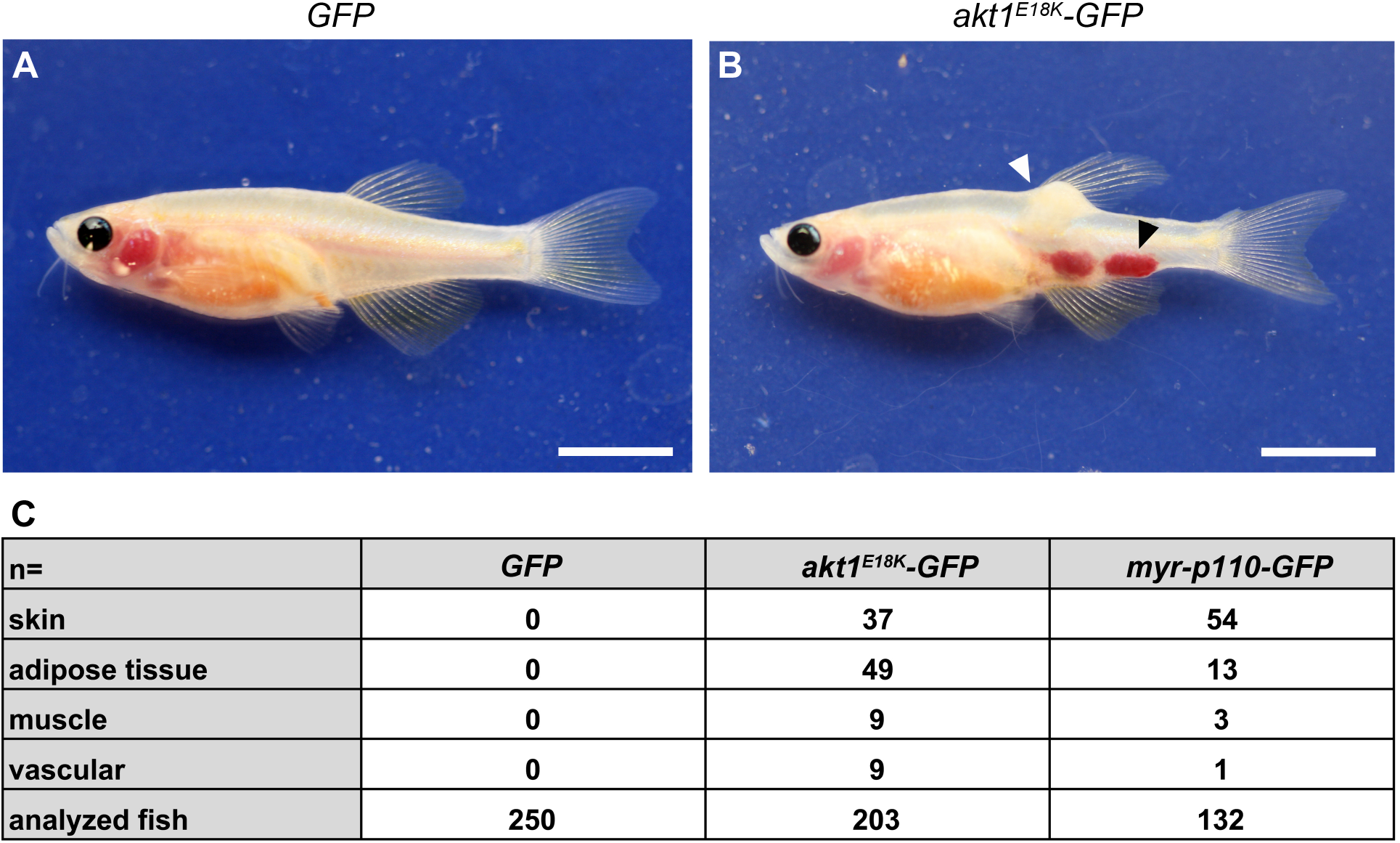
Zebrafish models for PROS and Proteus syndrome mimic patient phenotypes. (**A**) Representative adult *GFP* mosaic fish. (**B**) Example of an adult *akt1-GFP* mosaic fish with localized overgrowth of adipose tissue (white arrowhead) and vasculature (black arrowhead). **C**) Quantification of externally visible overgrowth phenotypes in 3-6 months old fish. Scale bars (**A, B**) 0.5 cm.

